# ICEs are the main reservoirs of the ciprofloxacin-modifying *crpP* gene in *Pseudomonas aeruginosa*

**DOI:** 10.1101/2020.03.13.991208

**Authors:** João Botelho, Filipa Grosso, Luísa Peixe

## Abstract

The ciprofloxacin-modifying *crpP* gene was recently identified in a plasmid isolated from a clinical *Pseudomonas aeruginosa* clinical isolate. Homologues of this gene were also identified in *Escherichia coli, Klebsiella pneumoniae* and *Acinetobacter baumannii*. We set out to explore the mobile genetic elements involved in the acquisition and spread of this gene in publicly available and complete genomes of *Pseudomonas*. The *crpP* gene was identified only in *P. aeruginosa*, in more than half of the complete chromosomes (61.9%, n=133/215) belonging to 52 sequence types, of which the high-risk clone ST111 was the most frequent. We identified 136 *crpP*-harboring ICEs, with 93.4% belonging to the mating-pair formation G (MPF_G_) family. The ICEs were integrated at the end of a tRNA^Lys^ gene and were all flanked by highly conserved 45-bp direct repeats. The core ICEome contains 26 genes (2.2% of all genes), which are present in 99% or more of the *crpP*-harboring ICEs. The most frequently encoded traits on these ICEs include replication, transcription, intracellular trafficking and cell motility. Our work reveals that ICEs are the main vectors promoting the dissemination of the ciprofloxacin-modifying *crpP* gene in *P. aeruginosa*.

**Author Notes:** All supporting data has been provided within the article or through supplementary data files. Supplementary material is available with the online version of this article.

**Impact Statement:** A high proportion of *Pseudomonas aeruginosa* clinical isolates are resistant to ciprofloxacin. Resistance to this antibiotic is often mediated by chromosomal mutations, but recently horizontally transferred genes have been identified. We assessed the repartition of the ciprofloxacin-modifying *crpP* gene among *Pseudomonas* genomes and we characterized the mobile elements associated with its acquisition. We found that this gene is prevalent in *P. aeruginosa* and frequently associated with integrative and conjugative elements (ICEs). Importantly, we also identified highly conserved direct repeats that can be used to accurately delimit *crpP*-carrying ICEs in *P. aeruginosa* genomes.

**Data Summary:** All the bacterial genomes scanned in this study have been deposited previously in the National Center for Biotechnology Information genome database and are listed on the supplementary tables. The newick files used to create the trees in Figures 1 and 4 are deposited on figshare at https://figshare.com/projects/ICEs_are_the_main_reservoirs_of_the_ciprofloxacin-modifying_crpP_gene_in_Pseudomonas_aeruginosa/79308.

## Introduction

*Pseudomonas aeruginosa* is a frequent cause of severe nosocomial infections and is one of the six ESKAPE pathogens [1,2]. Ciprofloxacin is an antibiotic of the fluoroquinolone class that is active against *P. aeruginosa* infections [3,4]. In this species, a high proportion of clinical isolates are resistant to ciprofloxacin [5,6]. Commonly reported mechanisms of ciprofloxacin resistance include mutations in DNA gyrase and topoisomerase IV-encoding genes *gyrA, gyrB, parC* and *parE* and efflux pumps regulatory genes as *nalC* and *nfxB* [6–8].

**Figure 1.**
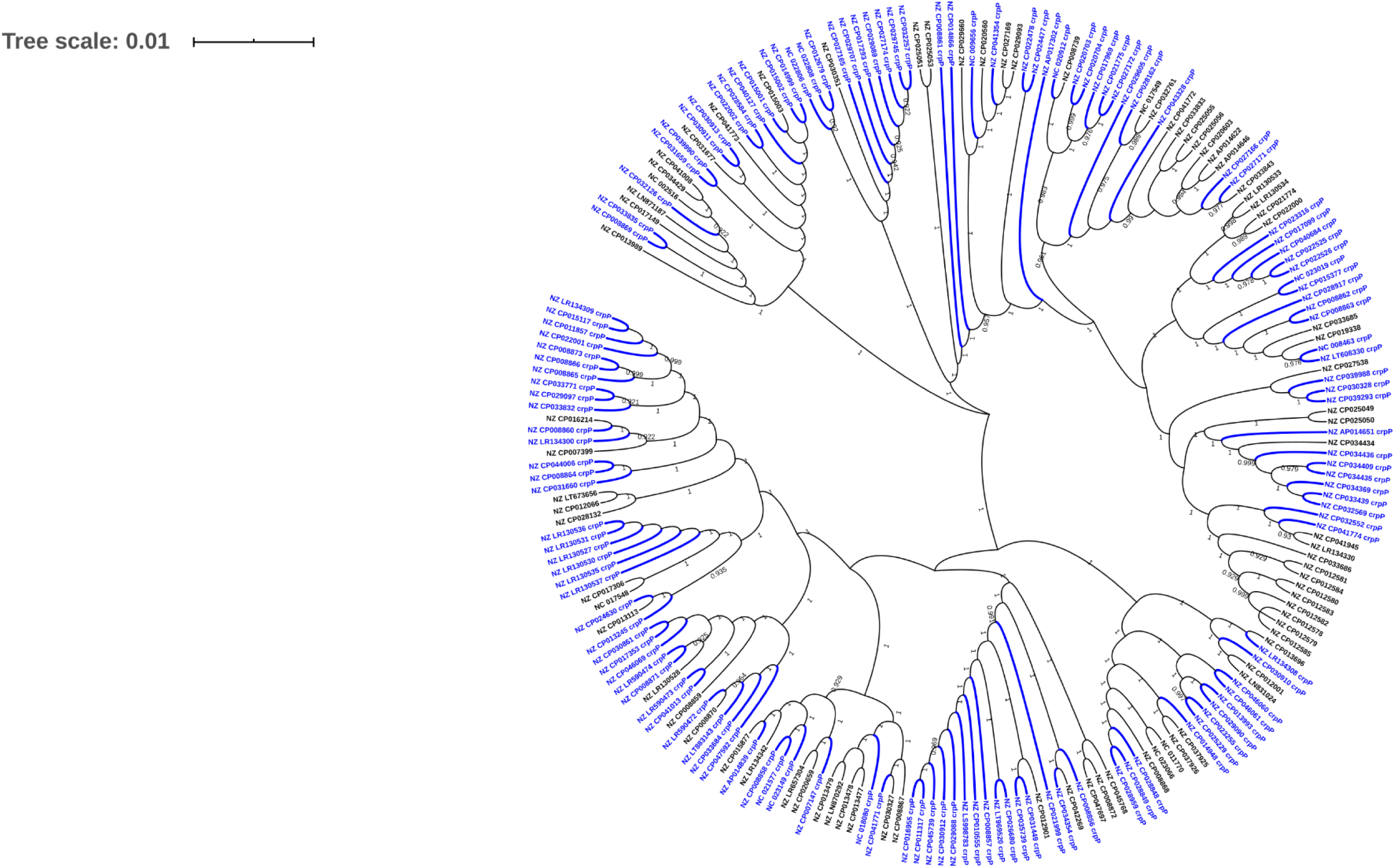
Approximately-maximum-likelihood phylogenetic tree base on the core genome alignment of 215 complete *P. aeruginosa* genomes. Branches and labels from *crpP*-positive hits are colored blue and the branches have twice the standard width. Only bootstrap values from 0.9 to 1 are displayed.

**Figure 2.**
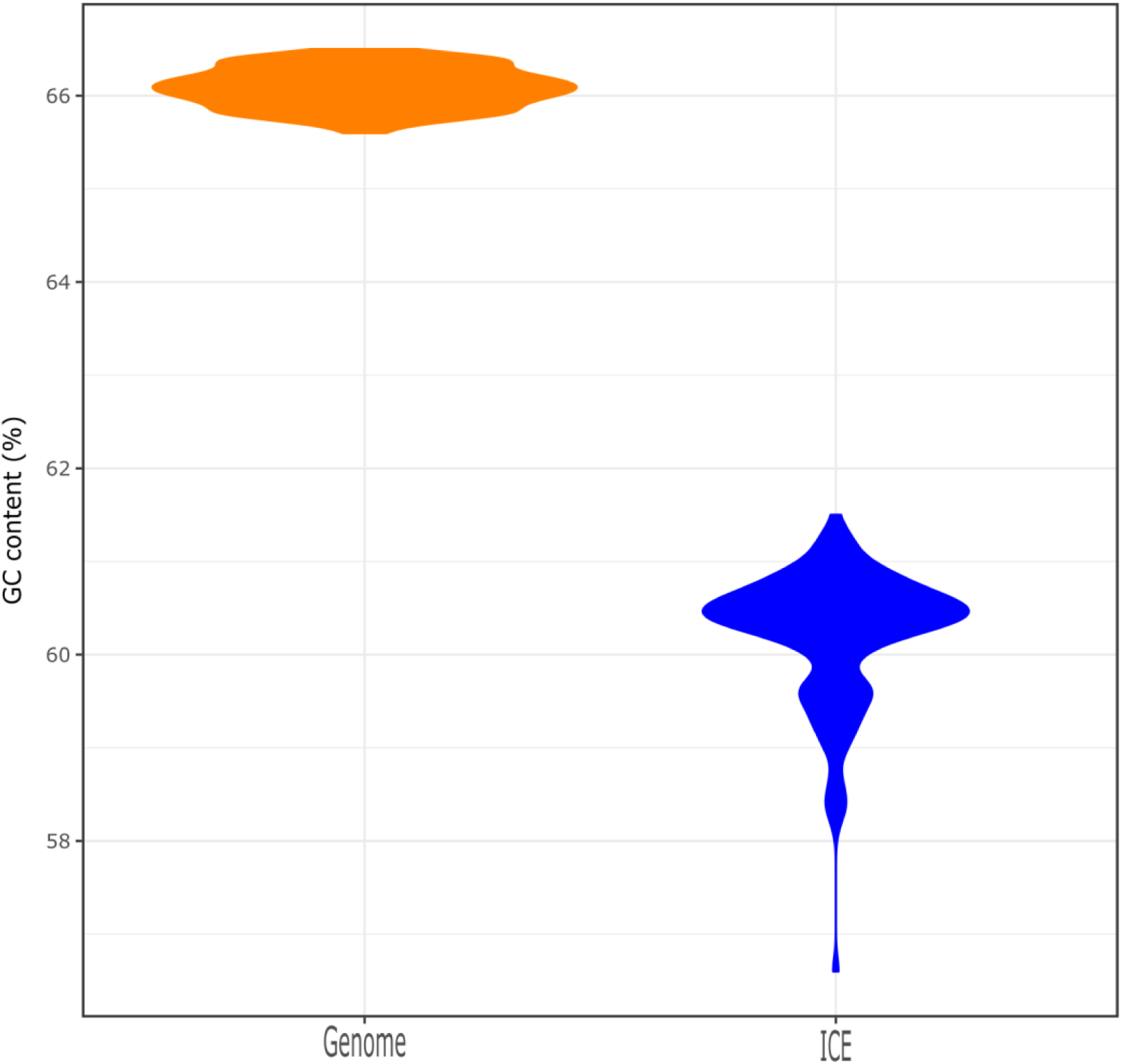
Distribution of the GC content of the chromosomes and ICEs. The violin plots were created using ggplot2.

**Figure 3.**
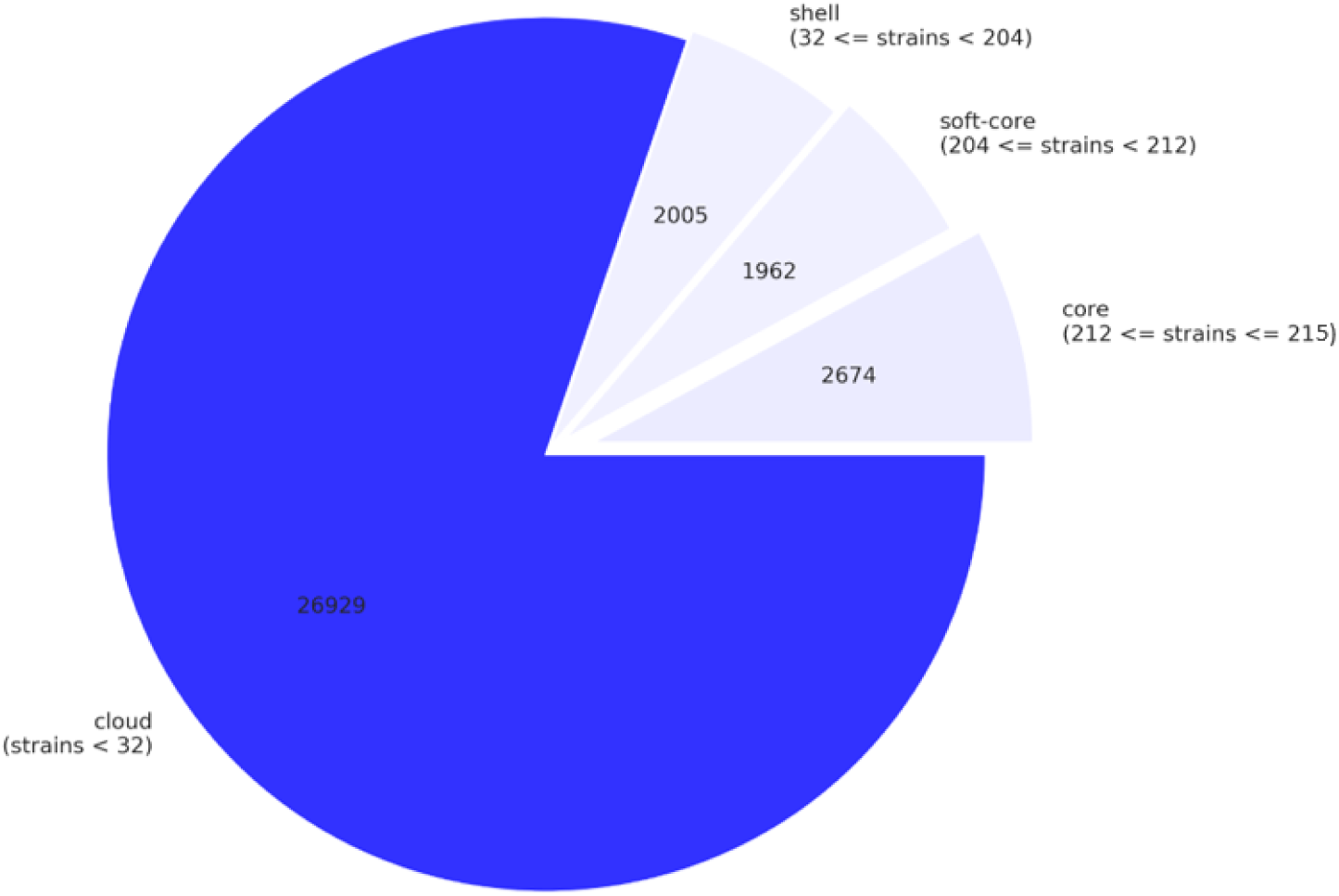
Pie chart of the breakdown of genes and the number of *P. aeruginosa* genomes they are present in. This figure was created using the contributed Python script roary_plots.py in https://github.com/sanger-pathogens/Roary/blob/master/contrib/roary_plots/roary_plots.py.

**Figure 4.**
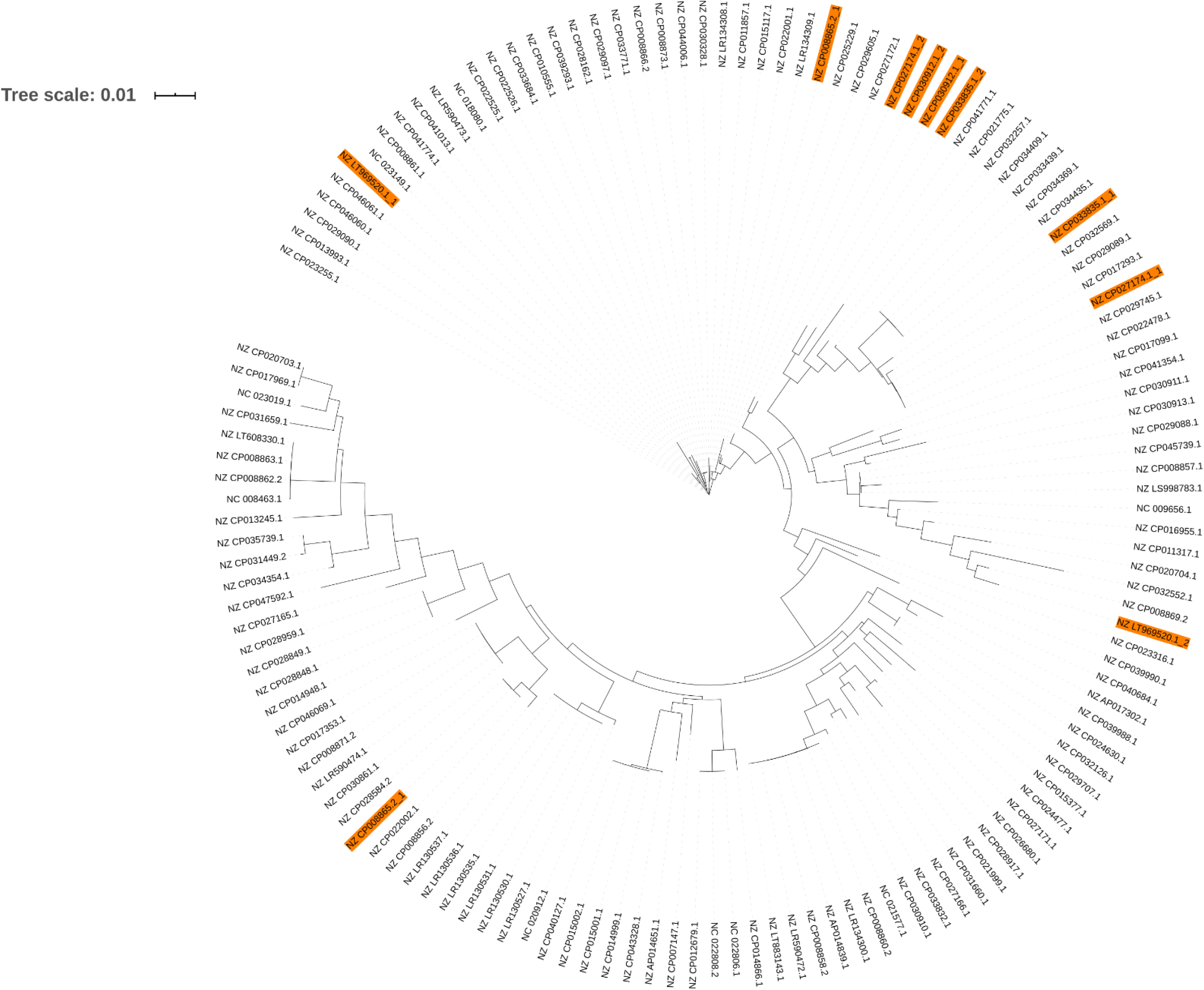
Approximately-maximum-likelihood phylogenetic tree base on the alignment of the 26 core genes identified in the *crpP*-harboring ICEs. Genome accession numbers containing more than one *crpP*-harboring ICE are highlighted in orange.

Besides chromosomal mutations, ciprofloxacin resistance can be mediated by horizontally transferred genes, such as the quinolone resistance *qnr* gene [9]. The ciprofloxacin-modifying *crpP* gene induces ATP-dependent phosphorylation of the antibiotic and was originally identified in a plasmid isolated from a *P. aeruginosa* clinical isolate [10]. This plasmid conferred resistance to ciprofloxacin when transferred to *P. aeruginosa* PAO1 strain. The authors also cloned the gene into a shuttle vector and the resulting recombinant plasmid conferred an increased minimum inhibitory concentration to the same antibiotic in *Escherichia coli*. Homologous *crpP* genes were also identified in *Acinetobacter baumannii* [11] and in *E. coli* and *Klebsiella pneumoniae* clinical isolates from Mexican hospitals and conferred decreased susceptibility to ciprofloxacin [12,13].

Integrative and conjugative elements (ICEs) are mobile elements involved in vertical and horizontal transmission of antibiotic resistance genes (ARGs) and other beneficial genes in bacterial communities [2,14,15]. ICEs can be classified into eight mating-pair formation (MPF) classes based on the classification of the secretion machinery involved in conjugation [16,17]. Recently, Ruiz verified that several *CrpP* proteins were encoded next to genes typically present in ICEs [11]. Building on this observation, we set out to trace and characterize the *crpP*-harboring ICEs present in all complete *Pseudomonas* genomes available on NCBI.

## Material and methods

### Pangenome and whole-genome analyses

All complete *Pseudomonas* genomes (n=577 chromosomes and 163 plasmids) were downloaded from NCBI’s Refseq library on the 01/02/2020 using ncbi-genome-download v 0.2.11 (https://github.com/kblin/ncbi-genome-download) (**Table S1**). We used mlst v 2.11 (https://github.com/tseemann/mlst) to scan the genomes against the PubMLST *Pseudomonas* typing schemes (https://pubmlst.org/) [18]. The genomes were annotated using prokka v1.14.5 (https://github.com/tseemann/prokka) [19]. We used the .gff files created by Prokka to calculate the *P. aeruginosa* pangenome in roary v3.13.0 (https://github.com/sanger-pathogens/Roary) [20]. The core genome and ICEome alignments created by Roary was used as input in fasttree v2.1.10 (http://www.microbesonline.org/fasttree/) [21] to create an approximately-maximum-likelihood phylogenetic tree using a generalized time-reversible (GTR) model of nucleotide evolution. We used the newick file created by FastTree to create a phylogenetic tree in iTOL (https://itol.embl.de/) [22]. We performed functional annotation based on orthology assignments (COGs) of protein files created by prokka using eggNOG-mapper v2 (http://eggnog-mapper.embl.de/) [23].

### Mining ICEs in complete genomes

We used the ICEfinder standalone version (http://202.120.12.136/ICEfinder/ICEfinder.html) and manual curation to trace putative ICEs. We searched for ARGs on extracted ICEs using amrfinder v 3.6.7 and default parameters (50% minimum coverage of the reference protein and 90% minimum identity) (https://github.com/ncbi/amr/wiki/AMRFinder-database) [24]. Positive ICE hits for CrpP-encoding genes were further characterized; we used fastANI v1.3 (https://github.com/ParBLiSS/FastANI) [25] to compute whole-genome average nucleotide identity of non-*P. aeruginosa crpP*-positive hits, antismash online tool (https://antismash.secondarymetabolites.org/#!/start) [26] to look for secondary metabolite biosynthesis gene clusters, BAGEL (http://bagel4.molgenrug.nl/) [27] to trace bacteriocins, macsyfinder v1.0.5 (https://github.com/gem-pasteur/macsyfinder) and the CRISPRCasFinder online tool (https://crisprcas.i2bc.paris-saclay.fr/) to detect CRISPRs and *cas* genes [28]. We used roary to infer the total *crpP*-harboring ICE content, here referred to as the ICEome.

## Results

### The *crpP* gene is widespread in *P. aeruginosa*

The *CrpP*-encoding gene was identified in 23.1% of the chromosomes (n=133/577, including 131 *P. aeruginosa*, 1 *Pseudomonas fluorescens* and 1 *Pseudomonas* sp.) and 1 *P. aeruginosa* plasmid (accession number NZ_CP030914.1), which is different than the pUM505 plasmid reported by Chávez-Jacobo and colleagues [10]. However, we compared the ANI between non-*P. aeruginosa* hits (the *P. fluorescens* strain NCTC10783 and *Pseudomonas* sp. AK6U strains with accession numbers NZ_LR134300.1 and NZ_CP025229.1, respectively) and the *P. aeruginosa* DSM 50071 type strain (accession number NZ_CP012001.1) and we realized that these strains belong to the *P. aeruginosa* species, as the ANI value is above the 95% cutoff for species delineation (**Table S2**). All *crpP*-hits were only identified in *P. aeruginosa* complete genomes, as so from here on we focused our attention on *P. aeruginosa* genomes. Indeed, the percentage increases if we only consider the *P. aeruginosa* chromosomes (61.9%, n=133/215), meaning that the *crpP* gene is present in more than half of the *P. aeruginosa* complete genomes (**Figure 1**). The GC content of *crpP*-harboring chromosomes varies from 65.6 to 66.5% (**Figure 2**) and the sequence length from 6.3 to 7.5 Mb. We identified several clones (n=52 sequence types), of which the high-risk clone ST111 [29] was the most frequent (**Table S3**).

### The majority of *crpP*-harboring ICEs belong to the MPF_G_ family

A total of 316 putative ICEs were identified among the 133 chromosomes. Each chromosome carries at least one ICE. Most *crpP* genes present in the chromosome were associated with an ICE (with the exception of *P. aeruginosa* strains W60856 and B17932 with the accession numbers NZ_CP008864.2 and NZ_CP034436.1, respectively) with 43.0% of the total putative ICEs (n=136/316) harboring the ciprofloxacin-modifying gene. In *P. aeruginosa* RW109 strain (accession number NZ_LT969520.1, position 5629923-5820312 bp), we identified two *crpP*-harboring ICEs in tandem (**Table S3**). ICE size varied from 81.6 to 145.5 kb, and the GC content from 56.6 to 61.5% (**Figure 2**). Most of the ICEs (n=129/136) ICEs belong to the MPF_G_ family (**Table S4**). For the remaining 7, no MPF family could be determined.

### The *crpP*-carrying ICEs integrate into a specific hotspot

All *crpP*-carrying ICEs identified in this study were integrated at the end of a tRNA^Lys^ gene, that was found alone or in a tRNA cluster with tRNA^Pro^ and tRNA^Asn^. The only exception was observed in the two contiguous ICEs identified in *P. aeruginosa* RW109 strain, where the second ICE was integrated at the end of the first one. The tRNA^Lys^ genes shared the exact same sequence. All ICEs were flanked by 45-bp (5’-TGGTGGGTCGTGTAGGATTCGAACCTACGACCAATTGGTTAAAAG-3’) highly conserved direct repeats (DRs). The 136 integrases identified in this study, however, only shared 36.1% amino acid identity.

### The *P. aeruginosa* pangenome and the *crpP*-carrying ICEome

The total number of genes among the 215 *P. aeruginosa* complete genomes (pangenome) is 33570. The core genome (genes present in 99% or more of the genomes) contains 2674 genes (8.0% of all genes, **Figure 3** and **Figures S1**). We also identified 11241 unique genes, which are found in only one strain [30].

The total number of genes in the 136 *crpP*-harboring ICEs (ICEome) is 1193. The core ICEome contains 26 genes (2.2% of all genes). The soft core content (genes present in more or less than 95% and less than 99% of ICEs) includes 11 genes, 129 shell genes (between 15 and 95%) and 1027 cloud genes (between zero and 15%). We identified 451 unique genes among the ICEome. These values suggest that *crpP*-harboring ICEs are not clonal, based on the small number of core genes and high number of cloud genes. Phylogenetic analysis of the core ICEome alignment reveals that these ICEs are very diverse and the core genes only share 28.0% nucleotide identity (**Figure 4**).

### The *crpP*-carrying ICEs encode other beneficial genes

The most frequently encoded traits on *crpP*-harboring ICEome include replication, transcription, intracellular trafficking, and cell motility (**Figure 5**). However, nearly one-third of the proteins for which a COG category was attributed (30.7%, n=211/687) encode for an unknown function (**Table S5**).

**Figure 5.**
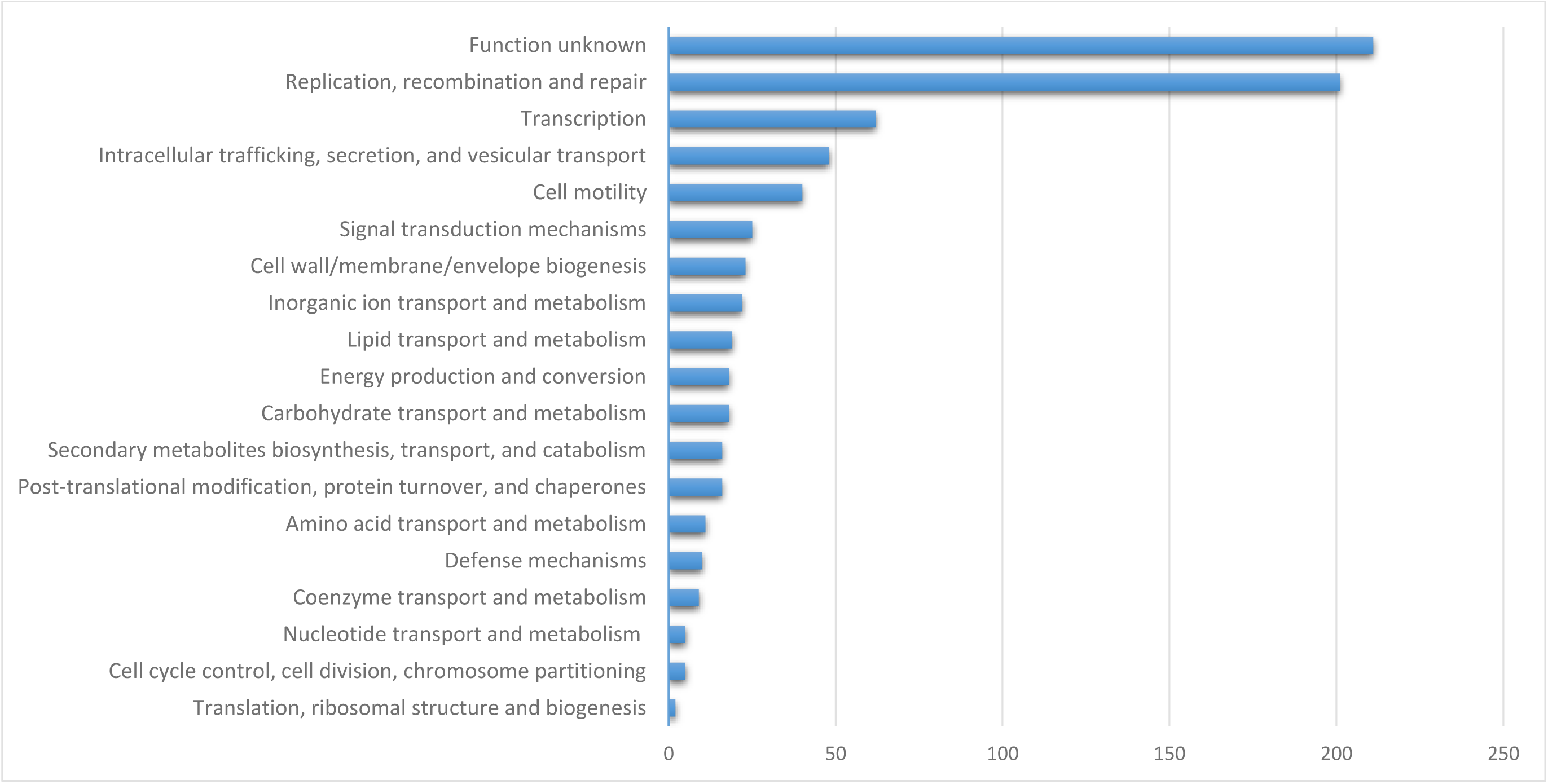
Functional annotation of proteins encoded by the *crpP*-harboring ICEome.

The *crpP* gene encoded for proteins ranging from 65 to 68 amino acid and is present in a single copy, with no flanking mobile units as integrons, insertion sequences and/or transposons or found on the close vicinity of the gene. The 136 CrpP proteins here identified share 19.7% amino acid identity. We also observed a lack of co-resident antibiotic resistance genes (ARGs) within the *crpP*-harboring ICEs. Seven ICEs carry a type I-C CRISPR-Cas system and ten harbor pyocin S5-encoding genes (**Table S3**). The bacteriocins here identified are highly identical, sharing 95% amino acid identity. Besides bacteriocins, no secondary metabolites were identified within the extracted ICEs.

## Discussion

Our results show that ICEs are the main drivers for the spread of the ciprofloxacin-modifying *crpP* gene in *P. aeruginosa*. A selective pressure exerted by the use of ciprofloxacin to treat *P. aeruginosa* infections may have promoted the dissemination of *crpP*-harboring ICEs among several clones.

Curiously, and in opposition to several ARGs, no integrons, insertion sequences or transposons were found on the vicinity of the *crpP* gene. Bacteriocins produced by bacteria are important for niche competition. The pyocin S5 possesses a bactericidal activity against clinical *P. aeruginosa* isolates [31,32]. The presence of these bacteriocins among some *crpP*-harboring ICEs can help to eliminate possible competitors and indirectly enhance ICE maintenance.

As expected for foreign DNA, the GC content of the ICEs is lower than that from the chromosomes. Still, given the high GC content of these mobile elements and the biological barrier for gene acquisition between donor-recipient pairs having >5% difference in GC values [33], we suspect that the ancestor of the *crpP* gene derives from GC-rich taxa.

We identified highly conserved DRs that can be used to accurately delimit *crpP*-carrying ICEs in *Pseudomonas*. Interestingly, the exact 45-bp sequence was found on the 2 ICEs from *P. fluorescens* and the *Pseudomonas* sp. AK6U strain. Also, even though the *crpP* gene was first reported in a plasmid in *P. aeruginosa* [10], we found a perfect match for the 45-bp DR next to an integrase-encoding gene. We only found one match for the DR in this plasmid, meaning that the ICE is degenerated and most likely depends on the plasmid to be mobilized. The same was observed for the *P. aeruginosa crpP*-carrying plasmid detected in this study (accession number NZ_CP030914.1).

The *crpP*-carrying ICEs were traced in several clones, taking advantage of the highly conserved attachment site in the tRNA^Lys^ gene. Interestingly, we found in a previous study that MPF_G_ class ICEs carrying specific ARGs were integrated at the end of tRNA^Gly^ genes [15]. In fact, this class of ICEs is frequently integrated at the end of a tRNA gene [34]. This behavior suggests selection for the maintenance of these non-coding integration sites, and seamless site-specific integration of ICEs in the chromosome will incur in a lower burden to the recipient cell and therefore increase its fitness [2,35]. As so, conferring decreased susceptibility or low levels of ciprofloxacin resistance without incurring in fitness cost can help to explain the high frequency of *crpP*-carrying ICEs observed in this study. Future studies should explore the possible synergistic effects between commonly reported mutations leading to ciprofloxacin resistance and the presence of *crpP*-harboring ICEs.

One of the caveats of our study is that we restricted the analyses to complete genomes. Given that the majority of the assemblies present in public databases as NCBI correspond to draft genomes, we probably missed the identification of several *crpP*-carrying ICEs among *P. aeruginosa*. However, mobile elements as ICEs tend to be fragmented in genomes sequenced with short-reads technologies, due to the presence of repetitive regions. Also, the identification of ICE fragments within the draft genome would require similar ICE sequences to use as a reference, which would then bias the study. Sequencing bacteria with long-read approaches is increasing, which will improve the accurate delimitation of an ever-growing number of mobile elements.

## Conclusions

Our work demonstrates that ICEs are the main promoters involved in the dissemination of the ciprofloxacin-modifying *crpP* gene among clonally-unrelated *P. aeruginosa* strains. We also identified highly conserved 45-bp DRs than can be used to accurately trace *crpP*-harboring ICEs on *P. aeruginosa* genomes. Future studies should explore the mobile elements involved in the acquisition of the *crpP* homologues identified in species other than *P. aeruginosa* and the possible synergistic effects between commonly reported mutations leading to ciprofloxacin resistance and the presence of *crpP*-carrying ICEs.

## Data bibliography

The DNA sequences used in this study are available in NCBI, under the accession numbers provided in supplementary tables S1-S5.

## Author contributions

J. B., F. G. and L. P. conceptualized the project. J. B. ran the analyses and examined data.

J. B. wrote the manuscript. J. B., F. G. and L. P. edited and revised the manuscript. All authors read, commented on and approved the final manuscript.

## Abbreviations

ARG: antibiotic resistance gene
CrpP: ciprofloxacin resistance protein
ICE: integrative and conjugative element
MPF: mating pair formation
NCBI: National Center for Biotechnology Information.

## References

1 Boucher, H.W. et al. (2009) Bad Bugs, No Drugs: No ESKAPE! An Update from the Infectious Diseases Society of America. Clin. Infect. Dis. 48, 1–12

2 Botelho, J. et al. (2019) Antibiotic resistance in *Pseudomonas aeruginosa* –Mechanisms, epidemiology and evolution. Drug Resist. Updat. 44, 100640

3 Paulsson, M. et al. (2017) Antimicrobial combination treatment including ciprofloxacin decreased the mortality rate of *Pseudomonas aeruginosa* bacteraemia: a retrospective cohort study. Eur. J. Clin. Microbiol. Infect. Dis. 36, 1187–1196

4 Klodzinska, S.N. et al. (2016) Inhalable antimicrobials for treatment of bacterial biofilm-associated sinusitis in cystic fibrosis patients: Challenges and drug delivery approaches. Int. J. Mol. Sci. 17, 1688

5 Pitt, T.L. et al. (2003) Survey of resistance of *Pseudomonas aeruginosa* from UK patients with cystic fibrosis to six commonly prescribed antimicrobial agents. Thorax, 58, 794–796

6 Rehman, A. et al. (2019) Mechanisms of ciprofloxacin resistance in *Pseudomonas aeruginosa*: New approaches to an old problem. J. Med. Microbiol. 68, 1–10

7 Bruchmann, S. et al. (2013) Quantitative contributions of target alteration and decreased drug accumulation to *Pseudomonas aeruginosa* fluoroquinolone resistance. Antimicrob. Agents Chemother. 57, 1361–8

8 Cantón, R. and Ruiz-Garbajosa, P. (2011) Co-resistance: an opportunity for the bacteria and resistance genes. Curr. Opin. Pharmacol. 11, 477–485

9 Belotti, P.T. et al. (2015) Description of an original integron encompassing *bla*VIM-2, *qnrVC1* and genes encoding bacterial group II intron proteins in *Pseudomonas aeruginosa*. J. Antimicrob. Chemother. 70, 2237–2240

10 Chávez-Jacobo, V.M. et al. (2018) CrpP is a novel ciprofloxacin-modifying enzyme encoded by the *Pseudomonas aeruginosa* pUM505 plasmid. Antimicrob. Agents Chemother. 62, e02629–17

11 Ruiz, J. (2019) CrpP, a passenger or a hidden stowaway in the *Pseudomonas aeruginosa* genome? J. Antimicrob. Chemother. 74, 3397–3399

12 Chávez-Jacobo, V.M. et al. (2019) Prevalence of the *crpP* gene conferring decreased ciprofloxacin susceptibility in enterobacterial clinical isolates from Mexican hospitals. J. Antimicrob. Chemother. 74, 1253–1259

13 Pablo, J. et al. (2020) Identification of Essential Residues for Ciprofloxacin Resistance of Ciprofloxacin-• Modifying Enzyme (CrpP) of pUM505. Microbiology DOI: 10.1099/mic.0.000889

14 Partridge, S.R. et al. (2018) Mobile Genetic Elements Associated with Antimicrobial Resistance. Clin. Microbiol. Rev. 31, e00088–17

15 Botelho, J. et al. (2018) Carbapenemases on the move: it’s good to be on ICEs. Mob. DNA 9, 37

16 Guglielmini, J. et al. (2011) The Repertoire of ICE in Prokaryotes Underscores the Unity, Diversity, and Ubiquity of Conjugation. PLoS Genet. 7, e1002222

17 Guglielmini, J. et al. (2014) Key components of the eight classes of type IV secretion systems involved in bacterial conjugation or protein secretion. Nucleic Acids Res. 42, 5715–27

18 Jolley, K.A. and Maiden, M.C.J. (2010) BIGSdb: Scalable analysis of bacterial genome variation at the population level. BMC Bioinformatics 11, 595

19 Seemann, T. (2014) Prokka: rapid prokaryotic genome annotation. Bioinformatics 30, 2068–2069

20 Page, A.J. et al. (2015) Roary: Rapid Large-Scale Prokaryote Pan Genome Analysis. Bioinformatics 31, 3691–3

21 Price, M.N. et al (2009) FastTree: Computing Large Minimum Evolution Trees with Profiles instead of a Distance Matrix. Mol. Biol. Evol. 26, 1641–50

22 Letunic, I. and Bork, P. (2019) Interactive Tree Of Life (iTOL) v4: recent updates and new developments. Nucleic Acids Res. 47, W256–W259

23 Huerta-Cepas, J. et al. (2019) eggNOG 5.0: a hierarchical, functionally and phylogenetically annotated orthology resource based on 5090 organisms and 2502 viruses. Nucleic Acids Res 47, D309–D314

24 Feldgarden, M. et al. (2019) Validating the AMRFinder Tool and Resistance Gene Database by Using Antimicrobial Resistance Genotype-Phenotype Correlations in a Collection of Isolates. Antimicrob. Agents Chemother. 63,

25 Jain, C. et al. (2018) High throughput ANI analysis of 90K prokaryotic genomes reveals clear species boundaries. Nat. Commun. 9, 5114

26 Blin, K. et al. (2019) antiSMASH 5.0: updates to the secondary metabolite genome mining pipeline. Nucleic Acids Res 47, W81–W87

27 van Heel, A.J. et al. (2018) BAGEL4: a user-friendly web server to thoroughly mine RiPPs and bacteriocins. Nucleic Acids Res. 46, W278–W281

28 Couvin, D. et al. (2018) CRISPRCasFinder, an update of CRISRFinder, includes a portable version, enhanced performance and integrates search for Cas proteins. Nucleic Acids Res. 46, W246–W251

29 Oliver, A. et al. (2015) The increasing threat of *Pseudomonas aeruginosa* high-risk clones. Drug Resist. Updat. 21–22, 41–59

30 Tettelin, H. et al. (2005) Genome analysis of multiple pathogenic isolates of *Streptococcus agalactiae*: Implications for the microbial “pan-genome.” Proc. Natl. Acad. Sci. U. S. A. 102, 13950–13955

31 Ling, H. et al. (2010) A predicted S-type pyocin shows a bactericidal activity against clinical *Pseudomonas aeruginosa* isolates through membrane damage. FEBS Lett. 584, 3354–3358

32 Elfarash, A. et al. (2014) Pore-forming pyocin S5 utilizes the FptA ferripyochelin receptor to kill *Pseudomonas aeruginosa*. Microbiol. (United Kingdom) 160, 261–269

33 Popa, O. et al. (2011) Directed networks reveal genomic barriers and DNA repair bypasses to lateral gene transfer among prokaryotes. Genome Res. 21, 599–609

34 Cury, J. et al. (2017) Integrative and conjugative elements and their hosts: composition, distribution and organization. Nucleic Acids Res. 45, 8943–8956

35 Touchon, M. et al. (2014) The chromosomal accommodation and domestication of mobile genetic elements. Curr. Opin. Microbiol. 22, 22–9

