## Supplementary material for "ICEs are the main reservoirs of the ciprofloxacin-modifying *crpP* gene in *Pseudomonas aeruginosa*": Figure S1

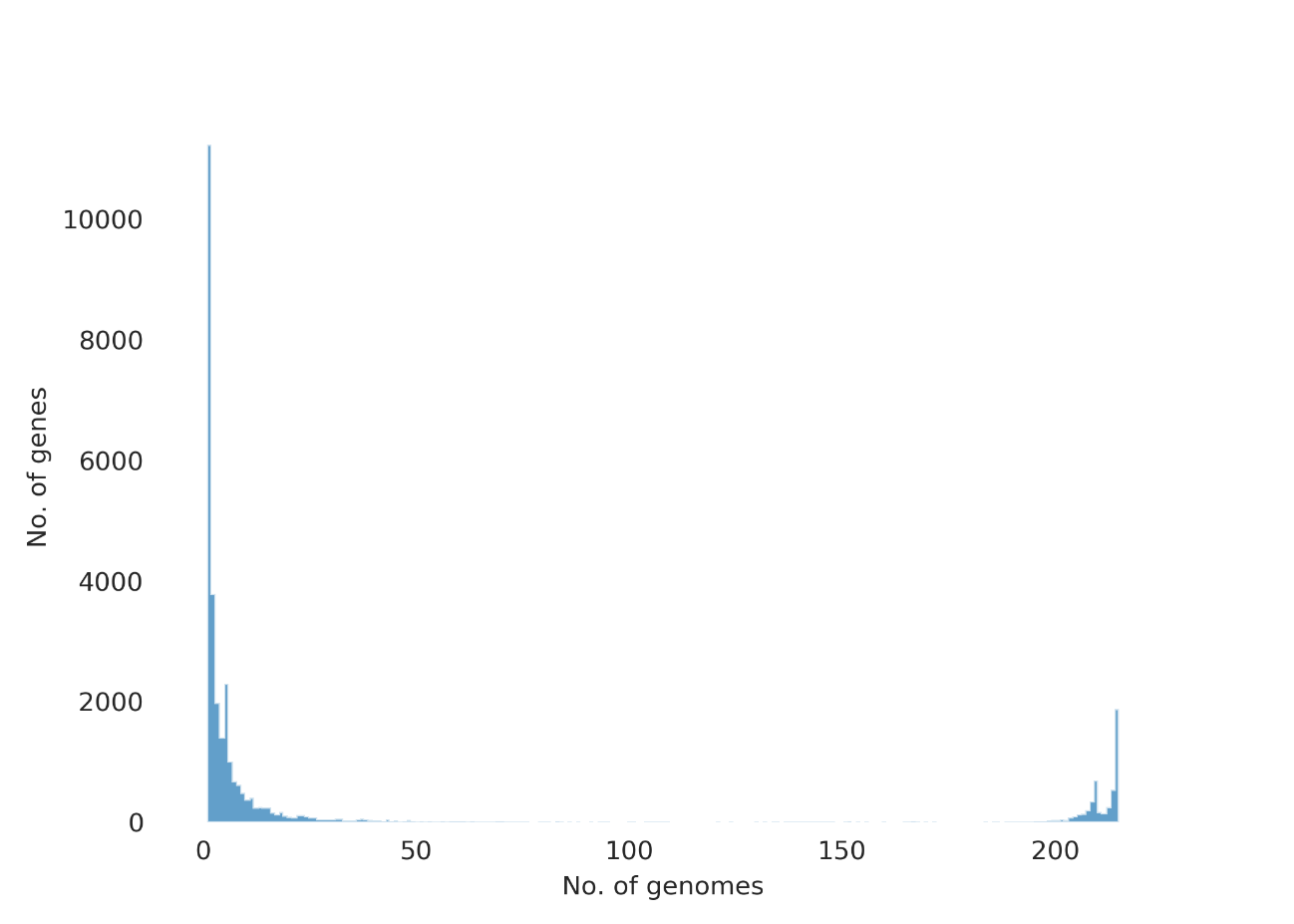

**Figure S1** - Graph with the frequency of genes versus the 215 *P. aeruginosa* complete genomes used in this study. This figure was created using the contributed Python script roary_plots.py in <https://github.com/sanger-pathogens/Roary/blob/master/contrib/roary_plots/roary_plots.py>.
